# Effects of Primed Adipose Mesenchymal Stem Cell-Derived Exosomes on Immunomodulation in Behcet Uveitis

**DOI:** 10.1101/2024.11.24.625042

**Authors:** Merve Gozel, Dilara Aydemir, Humeyra Nur Kaleli, Feray Cosar, Umit Yasar Guleser, Eda Kusan Unlu, Cem Kesim, Billur Sezgin, Afsun Sahin, Didar Ucar, Gulen Hatemi, Murat Hasanreisoglu

## Abstract

**Purpose:** This study sought to investigate the potential role of exosomes of interferon-gamma (IFN-γ) primed (IFN+Exo) and non-primed (IFN-Exo) adipose-derived mesenchymal stem cells (AdMSCs) for the treatment of Behçet Disease (BD) uveitis (BU).

**Methods:** AdMSCs were isolated from adipose tissue. Characterization and multipotency analyses were performed. Exosomes were isolated from the media of AdMSCs and then characterized. Peripheral blood mononuclear cells (PBMC) were isolated from patients with BU and healthy individuals. AdMSCs were preincubated with or without IFN-γ for 48 h. PBMC of patients with BU and healthy controls separately cultured with exosomes for 72 h. After the culture period, lymphocyte proliferation, viability, and apoptosis were carried out via flow cytometry. The expression of interleukin (IL)-10, IL-17, transforming growth factor (TGF)-β, and interferon (IFN)-γ levels were measured by real-time polymerase chain reaction (RT-PCR).

**Results:** IFN+Exo promoted lymphocyte apoptosis in patients with BU. IFN+Exo suppressed cell viability of T lymphocytes of BU. Exosomes alone did not affect on T lymphocyte proliferation. Additionally, exosomes increased anti-inflammatory cytokine levels and reduced pro-inflammatory cytokine levels of T lymphocytes in patients with BU.

**Conclusion:** This study’s findings can open a new pathway in MSCs/exosome therapy in BU.

## 1 Introduction

Behçet’s disease (BD) is a chronic, multisystemic inflammatory disorder characterized by relapsing inflammation of unknown etiology [1]. Classical clinical manifestations of BD include recurrent oral and genital ulcers, uveitis, and mucocutaneous lesions. BD can also affect the vessels of almost any system, including the gastrointestinal tract and nervous system [2]. BD is caused by immune system cells producing an excessive immunological response. Although genetic predisposition is generally shown as the most crucial cause of the disease, factors such as environmental, viral, or bacterial infections, as well as stress can trigger the onset of the disease [3–6]. Current literature also suggests that a Th1 and Th17 immune response predominates BD pathogenesis. High levels of cytokines related to Th1/Th17, including interleukin (IL)-2, interferon-gamma (IFN-γ), IL-6, IL-12, IL-17, and IL-23 have been reported in BD [7, 8]. BD uveitis is identified as panuveitis with non-granulomatous anterior segment inflammation and occlusive retinal vasculitis as its landmark features [9]. If untreated, BD uveitis has a poor prognosis due to inflammation and retinal damage, leading to irreversible vision loss [10]. Current treatment strategies that are used for BD uveitis include corticosteroids, conventional immunomodulators, and biological agents [11]. However, the lack of sufficient treatment response for current medications remains a major setback for BD management. Furthermore, these drugs are associated with several ocular and systemic side effects that cause treatment intolerance [12]. Therefore, there is a need for new treatment options that can effectively suppress ocular inflammation with the least potential adverse effects.

Stem cell therapy has recently found great potential for the treating chronic degenerative and inflammatory diseases. Mesenchymal stem cells (MSCs) are pluripotent stem cells extensively studied for their therapeutic potential. They can differentiate into other cells, migrate to damaged tissue, and exert immunomodulatory effects by inhibiting lymphocyte activation and proliferation, suppressing the secretion of inflammatory cytokines, reducing the function of antigen-presenting cells, and inducing regulatory T and B cells [13, 14].

Adipose-derived mesenchymal stem cells (AdMSCs) have been shown to exhibit immunomodulatory activity against T cells by expression higher IL-6 and transforming growth factor beta (TGF-β) than bone marrow MSC. AdMSCs allow easier access to stem cell therapy with higher efficiency and less ethical concerns [15]. Stem cells can affect damaged tissues with intracellular cytokine and signal interaction and increase their survival potential. A newly studied treatment strategy in autoimmune diseases introduces application of subcellular structures called exosomes to damaged tissue as an alternative of stem cell therapy [15, 16]. Exosomes are described as nano-sized extracellular vesicles released by fusion as various packages, including cell-specific proteins, lipids, and genetic material that stimulate cytokine/signal molecules [17]. They can provide therapeutic effects through the modulation of immunity, depending on their cellular origin and targets without having disadvantages of cellular therapy such as high immunogenicity, low repairability and difficulties to cross the barriers (blood-brain barrier, capillaries) [16, 18, 19].

In this study, we aimed to evaluate the effects of exosomes on changes in cell proliferation, apoptosis, and immunomodulation to determine whether the exosomes of AdMSCs primed with or without IFN-γ would reduce the abnormal inflammatory response from peripheral blood mononuclear cells (PBMC) of patients with BU.

## 2 Methods

### 2.1 Ethics Statements

This study adhered to national regulations and was approved by the Ethics Committee of the Koc University School of Medicine, Istanbul, Turkey (protocol no: 2020.447.IRB2.121, date of approval: 03.12.2020) and this study was performed in accordance with the Helsinki Declaration of 1964 and its later amendments. Study subjects (15 from BD with uveitis, 15 from healthy controls for blood samples and 8 adipose tissues from liposuction surgery) were recruited from Koç University Hospital, Ophthalmology Department, Rheumatology Department and Plastic, Reconstructive and Aesthetic Surgery Department. Written informed consent was obtained from all patients after an explanation of the nature and possible consequences of the study.

### 2.2 Isolation and culture of AdMSCs

Adipose tissues were collected in tubes containing Dulbecco’s phosphate-buffered saline (DPBS; Biowest, Nuaille, France), and penicillin/streptomycin (P/S; Biowest, Nuaille, France). The Lipoaspirates or abdominoplasty samples were washed extensively with equal volumes of DPBS, and the extracellular matrix then enzymatically digested at 37°C with 0.075% collagenase I (Chemicon®, Merck KgaA, Darmstadt, Germany). Enzyme activity neutralized with Dulbecco’s Modified Eagle Medium – low glucose (DMEM-LG; Biowest, Nuaille, France) containing 10% (v/v) Fetal Bovine Serum (FBS; Biowest, Nuaille, France) and 1% P/S, and samples centrifuged at 800 x g for 10 min. The pellets were washed with DPBS and filtered through a 70 μm nylon mesh (Cell Strainer, Sarstedt,

Nümbrecht, German). After filtration, cells were seeded into a T25 flask (TC-Flasche T25, Sarstedt, Nümbrecht, German) and incubated at 95% humidity at 37°C at 5% CO_2_. These cells were recognized as P0. After reaching 70-80% confluency, cells were trypsinized with Trypsin-EDTA (0.25%; Wisent Inc., Québec, Canada) and passaged into a T75 flask (recognized as P1). MSCs at P3 were used for characterization whereas cells at P4-P6 were used for isolation of exosomes (Fig. 1A).

**Figure 1.**
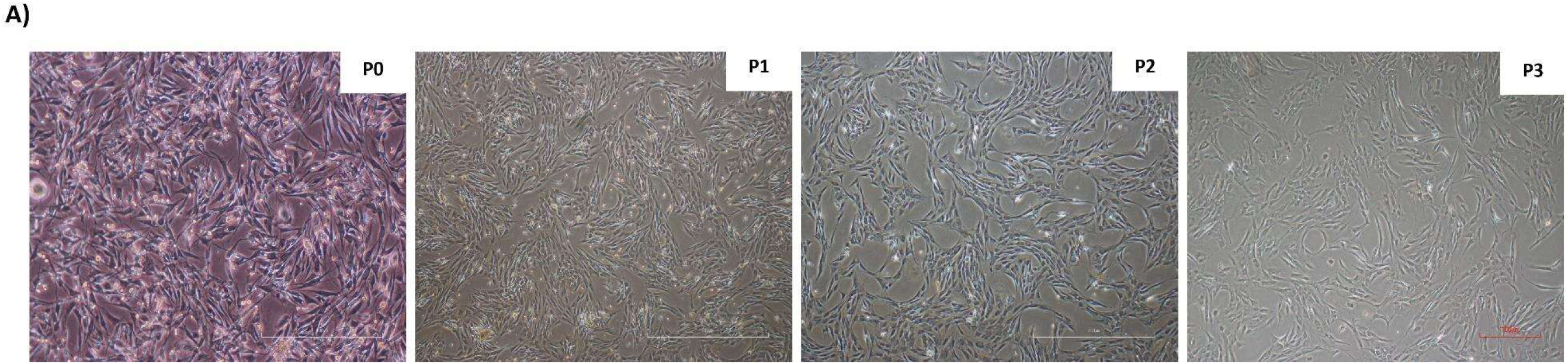
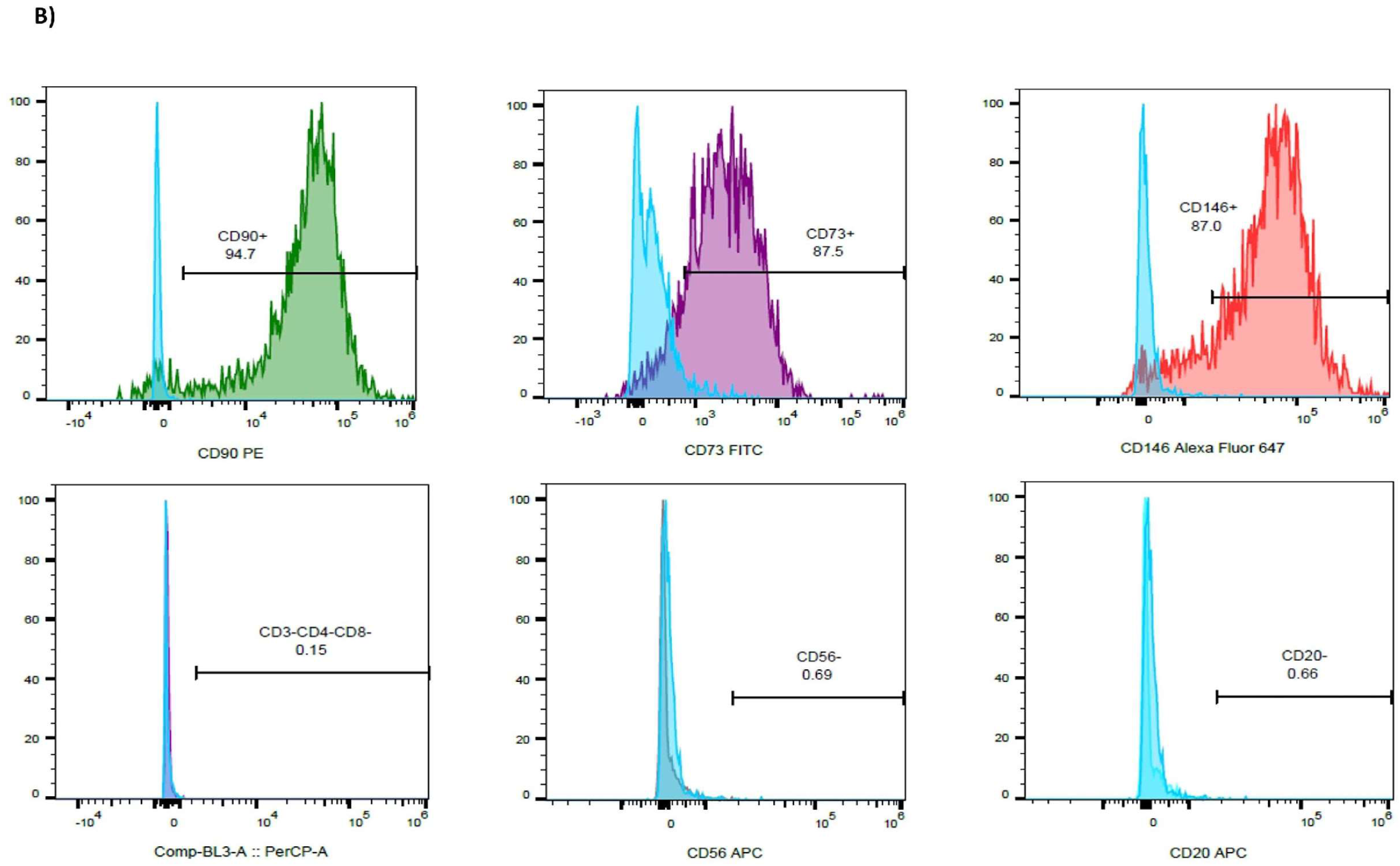
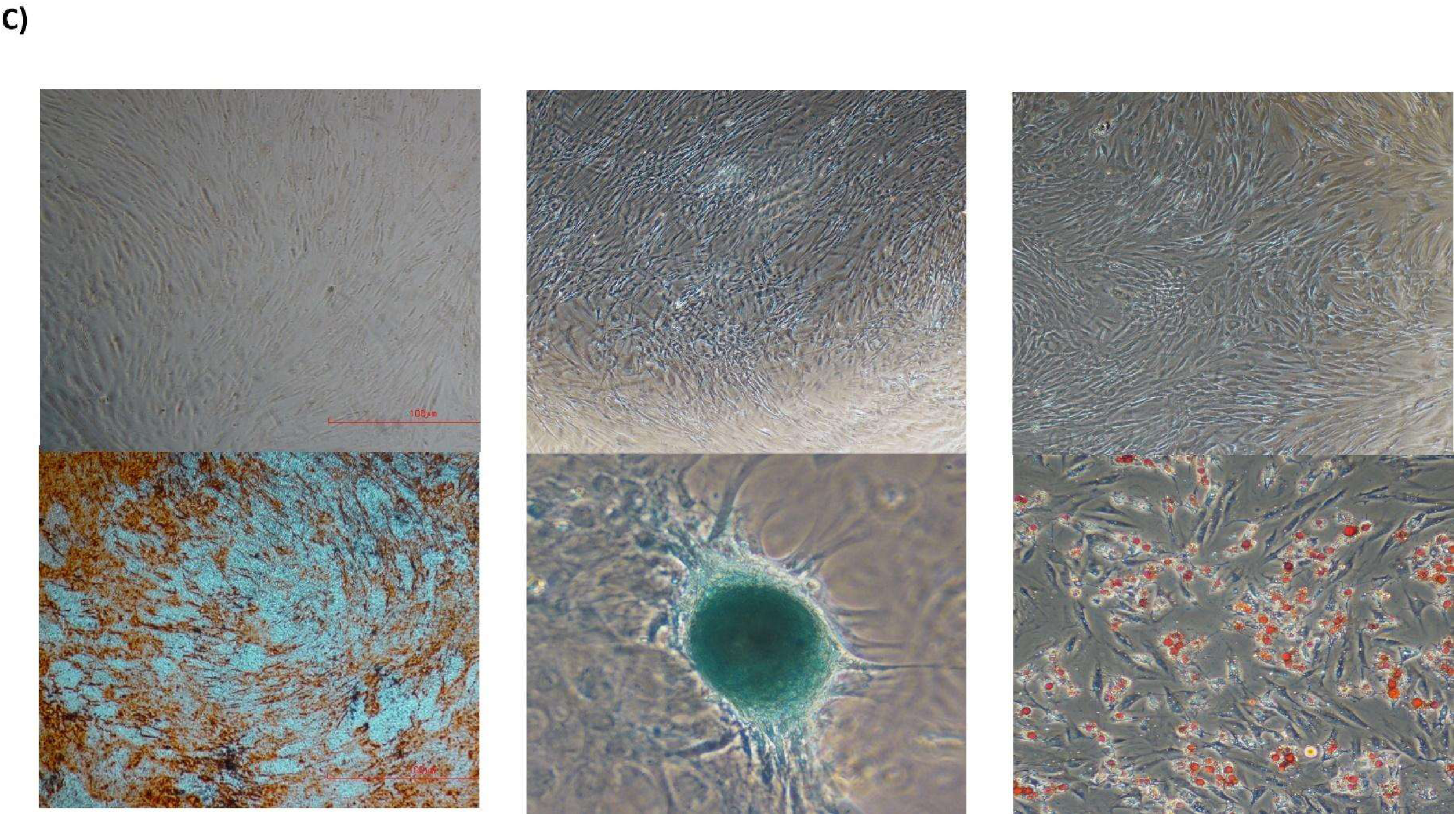
Characterization of Adipose-derived mesenchymal stem cells (AdMSCs). **(A)** Morphologic properties of AdMSCs. Representative brightfield image of AdMSCs at passages 0,1,2,3, respectively. Scale bar, 100 µm. Immunofluorescence imaging was captured with a 10x objective lens. **(B)** Expression of cell surface determinants using flow cytometry. The blue area represents unstained (negative control) cells, and the color areas depict marker expression. Cells at passage 3 were labeled with CD90, CD73, and CD146 as positive markers and CD3/4/8, CD56, and CD20 as negative markers. Values show the percentage of positive cells. **(C)** Assays for AdMSCs differentiation. Cells were cultured with lineage-specific media for 1 to 3 weeks to induce cell differentiation. Representative brightfield images of AdMSCs differentiated towards osteocytes (Alizarin red), chondrocytes (Alcian blue), and adipocytes (Oil Red O). The above represents negative controls. Scale bar, 100 µm. Immunofluorescence imaging was captured with a 10x objective lens.

### 2.3 Characterization of AdMSCs

AdMSCs were characterized using flow cytometry. After 70% confluency, the cells were detached using trypsin-EDTA (0.25%). Cell count was carried out, and the sample was homogenized in DPBS and incubated by adding 10 μl of specific Fluorescein isothiocyanate (FITC), and Phycoerythrin (PE) conjugated monoclonal antibodies with appropriate controls of the specified cell surface markers. Incubation was performed at room temperature (RT) in dark for 30 min. After incubation, a washing solution (DPBS containing 0.1% sodium azide) was added and centrifuged at 200 x g for 10 min, resuspended in a 500 μl washing solution, and analyzed with Attune NxT flow cytometer and its software package (FlowJo/V10, Ashland, OR, USA). In this study, we used antibodies against CD73 (FITC anti-human CD73; BioLegend, San Diego, California, USA), CD90 (PE anti-human CD90 (Thy1); BioLegend, San Diego, California, USA), and CD146 (Alexa Fluor® 647 anti-human CD146; BioLegend, San Diego, California, USA), as positive markers for MSCs and CD3/4/8 (CD4/8/3, BD Tritest™; BD Biosciences San Jose, CA) CD56 (NCAM16.2, APC CD56; BD Biosciences San Jose, CA), and CD20 (L27, APC CD20; BD Biosciences San Jose, CA) as negative markers for the hematopoietic cells (Fig. 1B).

### 2.4 Osteogenic, chondrogenic, and adipogenic differentiation assay

AdMSCs differentiation was performed when cell confluency was 80% in the T75 Flask (TC-Flasche T75, Sarstedt, Nümbrecht, Germany). The medium was discarded and washed with DPBS for 2 min. Then, the DPBS was removed, and Trypsin-EDTA (0.25%) was added and incubated for 4 min at 37°C. At the end of the incubation period, trypsin was inactivated with DMEM-LG supplemented with 10% FBS. Trypsinized cells were collected in falcon tubes and centrifuged at 500 x g for 5 min. The supernatant was discarded, the pellet was suspended in a 2 ml medium, and cells were counted and transferred to a new flask in an appropriate cell concentration. For osteogenic, chondrogenic, and adipogenic differentiation, cells were plated and grown until approximately 80% confluent, followed by osteogenic, chondrogenic, and adipogenic differentiation in a differentiation kit (Gibco, StemPro®, Grand Island, NY14072, USA). The cells were cultured for 2 weeks and then washed and fixed in 4% paraformaldehyde (4% PFA; Thermo Fisher Scientific, Waltham, Massachusetts, USA). The cells were stained with Alizarin Red S (Sigma-Aldrich, Darmstadt, Germany), which stains calcium deposits, Alcian Blue, which specifically stains the formulation of a chondrocyte, Oil Red O (Oil Red O Staining Solution, Sigma-Aldrich, Darmstadt, Germany), respectively and light microscopy images were captured at 10x (Fig. 1C).

### 2.5 Adipose-derived mesenchymal stem cells stimulation by IFN-γ

AdMSCs in the fourth passage were seeded in 145 cm^2^ plates (TC Dish 150; Sarstedt, Nümbrecht, Germany), 6×10^5^ cells cultured with culture medium. When the cells were in 60–80% confluence, the culture medium was replaced with a new medium with and without 20 ng/mL Interferon-gamma (IFN-γ; Bio Legend, San Diego, CA) for 48 h. Then the culture medium was replaced with serum-free media. After 72 h, the culture supernatant was collected for MSC-exosome isolation.

### 2.6 Exosome isolation

The ultracentrifugation method was used for exosome isolation. The conditioned serum-free media from human mesenchymal stem cells cultured for 72 h was pooled together for exosome isolation. The cellular debris was removed by centrifugation at 400 x g for 10 min, followed by centrifugation at 2000 x g for 10 min followed by centrifugation at 10,000 x g for 30 min to remove microvesicles. In the next step, DPBS was added to the ultracentrifuge tube. Then, 8 ml of conditioned media was loaded into DPBS solution, mixed well, and centrifuged at 100,000 x g at 4°C for 90 min using the Optima™ XPN-80 Ultracentrifuge from Beckman Coulter. The supernatant was discarded, and the exosome pellet was resuspended in 8ml of DPBS and washed by ultracentrifugation at 100,000 x g at 4 °C for 90 min to pellet down the exosomes. After this, the exosomes were resuspended in 50 μL DPBS and stored at – 80 °C for further analysis.

### 2.7 Exosomes Characterization

#### 2.7.1 Nanoparticle Tracking Analysis-based Exosome

The exosomes were diluted (1:10) in DPBS for nanoparticle tracking analysis (NTA) by NanoSight NS300 (NanoSight, Malvern Panalytical Ltd, Malvern, UK). Concentrations were found by pre-testing the ideal particle per frame value (20–100 particles/frame). A video of typically 60 seconds duration was taken, with a frame rate of 30 frames per second, and particle movement was analyzed by NTA software (v3.2, Malvern Panalytical, Malvern, United Kingdom).

#### 2.7.2 Scanning electron microscope imaging

The media was discarded, and the cells were washed with DPBS. 2% PFA was prepared from 16% PFA solution (Paraformaldehyde 16% Solution, EM Grade, Electron Microscopy Sciences, Hatfield, PA) in nanopure water. The cells were fixed using a 2% PFA solution and let air dry for 2 days. Scanning electron microscope (SEM) imaging was performed by ZEISS EVO LS15 Scanning electron microscope (Zeiss, Jena, Germany).

#### 2.7.3 Flow cytometry

50 μg exosomes were incubated with 5 μm aldehyde/sulfate latex beads (4% w/v, 4 µm, Invitrogen™, Massachusetts, United States) in a 100 μl final volume of DPBS and incubated for 15 min on a shaker at RT. Next, 200 μl 2% bovine serum albumin (BSA, Sigma Aldrich, Missouri, United States) in DPBS solution was added to the mixture and incubated for 2 h on a shaker. Then, 20 μL of 1 M glycine/DPBS was added and mixed gently to block the unbound sites of the latex beads and incubated for 30 min at RT. Exosomes/beads complex were washed twice with 800 μl cold DPBS by centrifugation at 2700 x g for 3 min. Exosomes were resuspended with 100 μl DPBS. The exosomes/beads complexes were incubated with primary anti-human antibodies against CD63 (anti-CD63 antibody [EPR5702], Abcam, Cambridge, MA, USA) overnight at 4°C. A negative control antibody reaction was performed using latex beads alone. Afterward, exosomes/beads complexes were washed with DPBS and incubated with secondary antibody for 30 min at RT. The labeled exosomes/beads complex were pelleted and washed twice as above with 1mL of DPBS. Finally, 100 μL pellets were resuspended in 400 μL of focusing fluid and subjected to flow cytometry (Attune NxT; Thermo Fisher Scientific).

#### 2.7.4 Western blotting

Western Blot was done by suspending the exosomes. The protein concentration of the exosome was measured by using a bicinchoninic acid assay (BCA) protein assay kit (Pierce, part of Thermo Fisher Scientific, Waltham, Massachusetts, USA). Exosomes were heated for 10 min at 70°C after mixing with the loading buffer. Equal volumes of exosomes were subjected to 4-12% SDS-PAGE in non-reducing conditions. Then, proteins were transferred to a polyvinylidene difluoride (PVDF) membrane (Bio-Rad Laboratories, Hercules, CA, USA). The blot was blocked in 5% non-fat skimmed milk (Blotting-Grade Blocker, Bio-Rad, USA) in 1X Tris-buffered saline with 0.1% Tween® 20 Detergent (TBS-T) solution followed by incubation in anti-CD63 (anti-CD63 antibody [EPR5702], Abcam, Cambridge, MA, USA), anti-Hsp70 [clone 1B5] (heat shock protein70, Enzo Life Sciences, Exeter, UK), anti-TSG101 (Anti-TSG101 antibody [EPR7130(B)], Abcam, Cambridge, MA, USA) and anti-calnexin (anti-calnexin primary antibody [EPR3633(2)], Abcam, Cambridge, MA, USA) overnight at 4°C. The membrane was washed and incubated with an HRP-conjugated anti-mouse IgG secondary antibody and developed using an ECL (Pierce™ ECL Plus-Chemiluminescent Substrates, ThermoFisher Scientific, Massachusetts, United States) imager using the ChemiDoc MP Imaging System (Bio-Rad Laboratories, Hercules, CA, USA).

### 2.8 Peripheral Blood Mononuclear Cells (PBMC) sample collection and cell isolation

Peripheral blood samples were purified from buffy coats of BU and healthy adult volunteer blood donors. The blood was transferred to 15 ml sterile falcon tubes in the UV cabinet, the sterile environment to isolate PBMC. An equal volume of DPBS was added and mixed well; then, 5 mL Ficoll-Paque plus solution (Pharmacia LKB, Uppsala, Sweden; density 1.077 g/L) was added per 10 mL blood. The sample was centrifuged for 20 min at 800 x g at RT without brake. The isolated PBMC was washed by adding excess DPBS and centrifuged for 10 min at 1000 x rpm at RT; the washing procedure was repeated. PBMC were resuspended in a cryopreservation medium consisting of 90% FBS and 10% dimethyl sulfoxide (DMSO, Sigma LifeScience, Riedstr, United Kingdom) and stored at – 80°C until use.

### 2.9 PBMC culture, stimulation, and exosome treatment

PBMC isolated from blood samples which taken from **patient with Behçet Disease uveitis (BU) and healthy control (HC) lymphocytes** were seeded in 96-well tissue culture plates at a density of 100,000 cells per well in RPMI Media (RPMI 1640, LONZA, Verviers, Belgium) containing 10% FBS and 1% P/S. After being stimulated with anti-CD3/anti-CD28 (Cdmix) (Dynabeads™ Human T-Activator CD3/28, Gibco, Vilnius, Lithuania), lymphocytes were cultured with 50 µg/ml **exosomes of IFN-γ primed (IFN+Exo) and non-primed (IFN-Exo) AdMSCs** for 72 h in a 37°C 5% CO2 incubator. Following this stage, cell viability and proliferation were assessed in the manner described below.

### 2.10 CFSE Assay

The proliferation of BU and HC lymphocytes were evaluated by labeling PBMC with 5 μM CFSE for 15 min at 37 °C. Then, the cells were washed with DPBS and centrifuged at 2000 x rpm for 5 min, and cells were resuspended in RPMI media containing 10% FBS and 1% P/S (culture media). PBMC were seeded in 96-well tissue culture plates at a density of 100,000 cells per well in culture media. CFSE-labeled lymphocytes were stimulated with Cdmix and were cultured with exosomes for 72 h. PBMC were analyzed for CFSE signaling by flow cytometry (CytoFLEX S, Beckman Coulter Life Sciences, California, United States).

### 2.11 Annexin V/PI Assay

Cell viability was quantified by staining for Annexin V (1 μg/ml) and Propidium Iodide (PI) (1 μg/ml) according to the manufacturer’s instructions (FITC Annexin V Apoptosis Detection Kit with PI, BioLegend, San Diego, CA, USA). The staining of Annexin V requires the usage of a binding buffer for Annexin V as a staining buffer. 100 μl of 1X staining buffer was added to the cells, and 2 μl of diluted FITC-Annexin V and 4 μl of PI solution were added to the cells. The stained cells were incubated at room temperature for 15 min in the dark. 300 μl Annexin V binding buffer was added to stained cells. All tubes were analyzed by flow cytometry.

### 2.12 Fas/Fas Ligand Assay

After the homogenized distribution of cells was ensured by pipetting into the wells; the homogenized liquid was transferred to Eppendorf tubes. DPBS was added to the tubes and centrifuged at 2000 x rpm for 5 min. The supernatant was removed, and 100 µl of DPBS was added to the tubes. 1 μl anti-human CD95 (FITC anti-human CD95 (Fas) Antibody, BioLegend, San Diego, CA) and 1 μl anti-human CD178 (PE anti-human CD178 (Fas-L) Antibody, BioLegend, San Diego, CA) were placed in each tube. The tubes were vortexed gently and left in the dark at room temperature for 15 min. At the end of the period, 400 µl of DPBS was added to each tube and centrifuged at 2000 x rpm for 5 min. The supernatant was discarded, 400 µl of DPBS was added, and after vortexing, they were measured by flow cytometry.

### 2.13 Quantitative real-time polymerase chain reaction (qRT-PCR)

To evaluate the effect of exosomes on Cdmix stimulated lymphocytes, the levels of mRNA of TGF-β, IFN-Ɣ, IL-17, and IL-10 were examined. Table S1. Provides a list of the primer sequences. Glyceraldehyde-3-Phosphate Dehydrogenase (GAPDH) served as an internal control. Following the manufacturer’s instructions, total ribonucleic acid extraction was carried out using the QuickRNA Microprep Kit (Zymoresearch, Irvine, CA, USA). The purity of the RNA was determined using the A260/280 and A260/230 ratios via spectrophotometer (NanoDrop 2000c, Thermo Scientific, USA). iScript™ cDNA Synthesis Kit (Bio-rad, 1708890) was used to reverse transcribe the RNA into complementary deoxyribonucleic acid (cDNA). Quantitative real-time PCR was carried out using 2 µl of 10-fold diluted cDNA in a final volume of 10 µl with the LightCycler 480 SYBR Green I Master 2X (Roche Diagnostics GmbH, Mannheim, Germany) following the manufacturer’s recommendations. The reaction steps are as follows: initial denaturation at 95°C for 10 min, followed by 40 cycles at 95°C for 15 sec and 60°C for 60 sec. Each process used two parallel samples intra-assay. The reactions were conducted using the Quantstudio 7 Real-Time PCR System (Applied Biosystems, CA, USA). The relative mRNA expression levels of the target genes were quantified using the 2-ΔΔCT technique.

### 2.14 Statistical analysis

The statistical analysis was done with the aid of GraphPad Prism 8.0 (GraphPad Software, La Jolla, CA, USA). One-way analysis of variance (ANOVA) with Tukey’s and Dunnett’s multiple comparisons was used for multi-group comparisons, and a two-tailed unpaired Student’s t-test was used for comparisons between two groups, and statistical significance was determined by P values less than 0.05.

## 3 Results

### 3.1 Isolation and characterization of AdMSCs

Under a light microscope (Nikon eclipse TS100, Inverted Routine Microscope, Marshall Scientific, Hampton, USA), adipogenic stem cells isolated from fat tissues appeared in culture flasks 3 days after plating. They looked like fibroblastic spindle-shaped cells and grew in monolayers in the culture flask after continued passaging (Fig. 1A). Fig. 1B shows the cell surface antigenic characteristics of the cultured AdMSCs at passage three by flow cytometry. The analyses revealed the expression of surface antigens, such as CD90, CD73, and CD146 were strongly positive, and CD3/4/8, CD56, and CD20 were negative. MSCs are defined as multipotent stem cells that should be able to differentiate into specific lineages including chondrocytes, osteoblasts, and adipocytes [20]. Therefore, to determine whether isolated MSCs could differentiate between these formations, the adipogenic, osteogenic and chondrogenic differentiation assays were performed. Alizarin staining clearly showed the formation of calcium oxalates. In addition, Alcian Blue Staining demonstrated the formation of chondrocytes (Fig. 1C). Intracellular lipid droplets staining using Oil Red O demonstrated the adipogenesis in MSCs (Fig. 1C). These observations were absent in the undifferentiated MSCs. These findings confirmed the characterization of cells as MSCs and showed the potential of MSC to differentiate into these lineages.

### 3.2 Exosome isolation and characterization

The exosome isolation of the target cell was successfully obtained through ultracentrifugation and centrifugation steps. The following methods were applied to evaluate the quality of exosomes; nanoparticle tracking analysis (NTA) to measure the size and concentration of exosomes, scanning electron microscopy (SEM) to observe the morphology of exosomes, Western blotting to detect exosome-related proteins, and flow cytometry to measure specific cell surface markers. NTA (Figs. 2A, 2B) and SEM (Fig. 2C) images proved exosome isolation in terms of shape and size. As shown in Fig. 2, exosomes with size between 30-400 nanometers (nm) were mostly captured. CD63 is a transmembrane protein that is enriched in the membrane of endosomes and lysosomes. As being transmembrane vesicles, exosomes are derived from endosomes and CD63 is commonly used as a specific marker for characterization of them. In flow cytometry analysis, it is found that 98.3% of AdMSC derived exosomes expressed CD63 protein (Fig. 2D). Tumor susceptibility gene 101 (TSG 101) and heat shock protein 70 (HSP 70) are other most used exosome markers along with CD63. To confirm isolated exosomes’ characteristics, expression of HSP 70 and TSG 101 were examined with western blotting (Fig. 2E). Exosomes are derived from endosomes, and they are devoid of ER specific proteins such as calnexin. Herein the expression level of calnexin was also evaluated in western blotting. Overall, isolated exosomes had spherical in shape with in diameter from 30 to 400 nm and they presented endosomal signaling markers as CD63, HSP70 and TSG 101.

**Figure 2.**
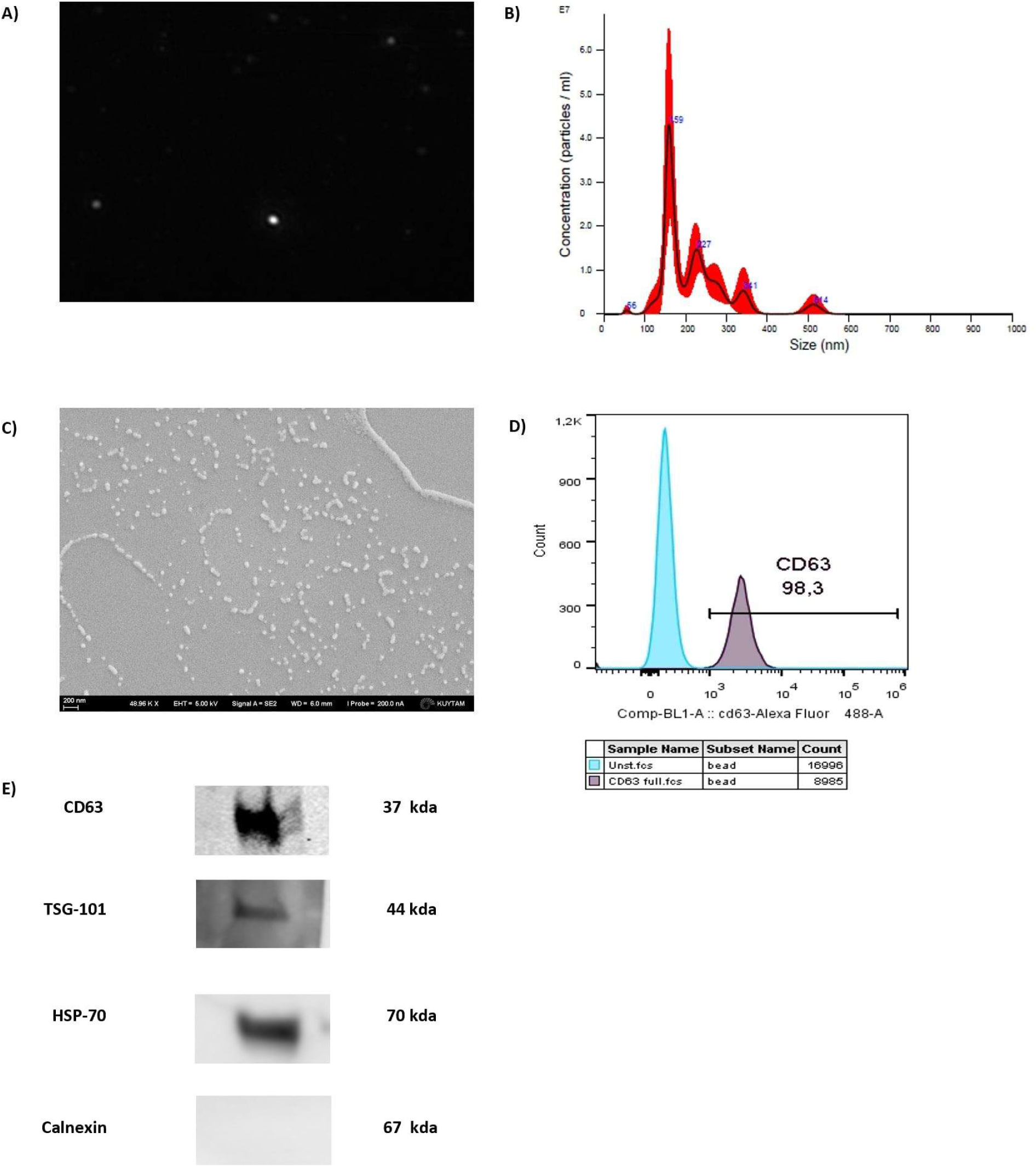
Characterization of exosomes. **(A, B)** Nanoparticle tracking analysis (NTA) analysis showed round particles with a size distribution of 30 to 400 nm. **(C)** Scanning electron microscopy (SEM) imaging showed a round morphology with a central depression, which is characteristic of exosomes. **(D)** Flow cytometry results showed high positivity for CD63 expression. The blue area represents unstained (negative control) cells, and the color area depicts marker expression. **(E)** Western blot results showed the presence of positive exosome markers CD63, HSP70 and TSG-101 as well as the absence negative exosome marker Calnexin in isolated exosomes.

### 3.3 Determination of cell proliferation of lymphocytes by CFSE

Previous studies show that MSC derived exosomes with a 50 µg/ml concentration was determined as the optimal exosomal protein concentration to induce apoptosis, inhibit proliferation, and prevent inflammation [21, 22]. Therefore, we continued the experiments with the proposed dose of the exosome. To investigate the effect of exosomes on cell proliferation of patients with BU and HC lymphocytes, CFSE-labeled lymphocytes were cultured with 50 µg/ml of exosomes for 72h. CFSE labeling is the method to determine cell proliferation by the rate of cell division. The dye is evenly distributed between daughter cells during cell division and the fluorescent yield after treatment condition is measured by flow cytometry. As shown in Fig. 3, neither the proliferation of HC lymphocytes nor that of BU lymphocytes was impacted by exosomes of IFN-γ primed (IFN+Exo) and non-primed (IFN-Exo) AdMSCs.

**Figure 3.**
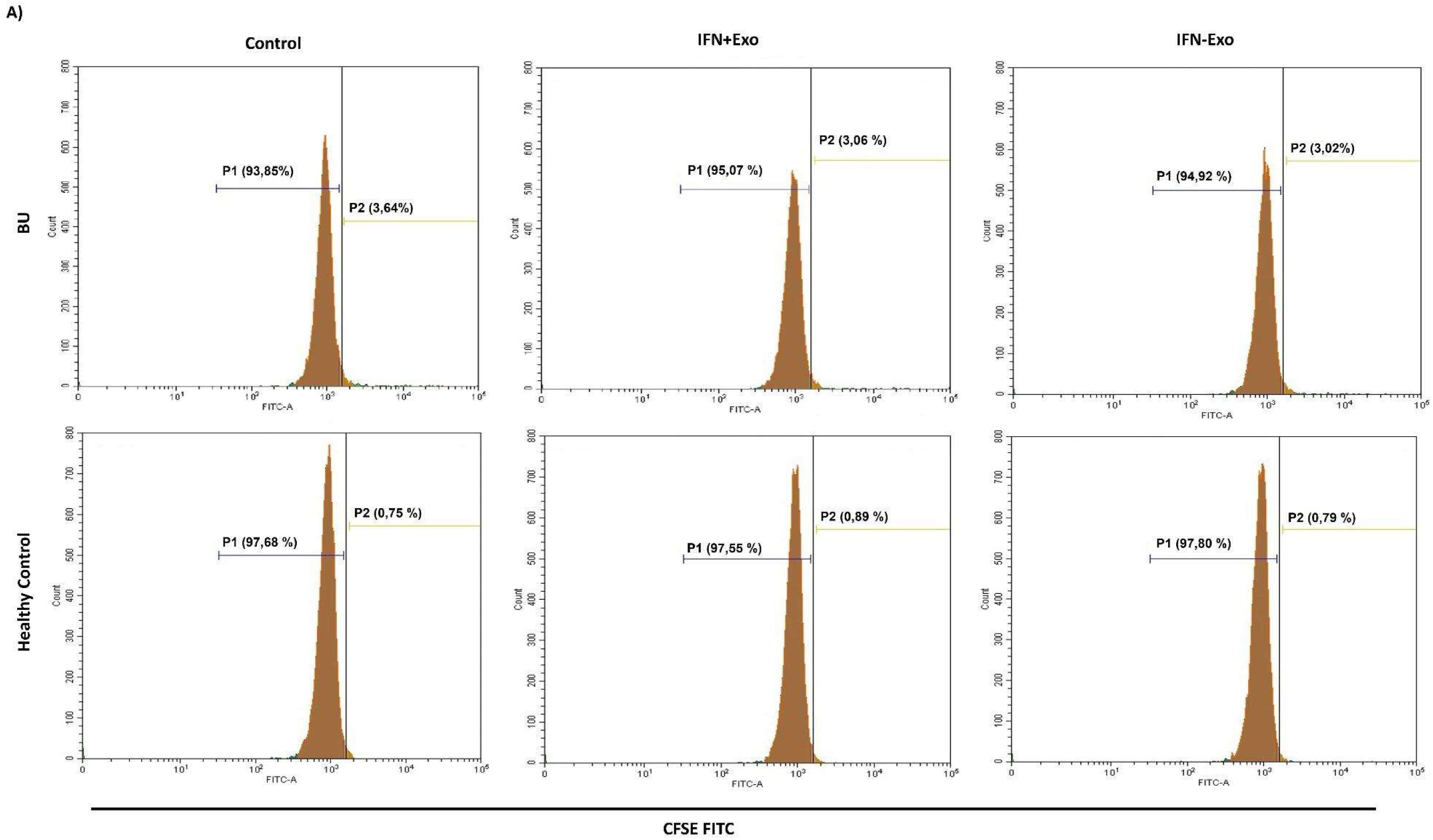
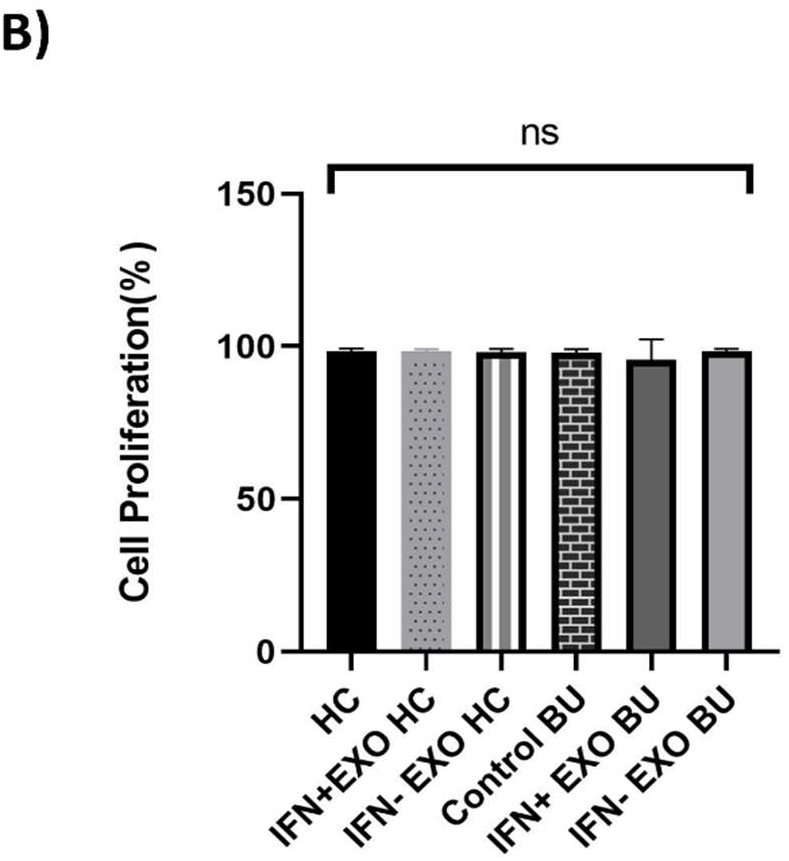
Inhibitory effect of exosomes on the proliferation of lymphocytes as detected by CFSE. **(A)** After 72h of culture, the effect of exosomes on the proliferation of lymphocytes is displayed by flow cytometry. P1 defines CFSE stained cells. IFN+/− Exo treatments did not change lymphocyte proliferation compared to control of healthy individuals. **(B)** Quantitative analysis of (A) (n=15). P > 0.05. Results are shown as mean ± SD. patients with BD uveitis *(BU), Exosomes of IFN-γ primed (IFN+Exo) and non-primed (IFN-Exo) AdMSCs, Healthy Controls (HC)*.

### 3.4 Detection of cell viability effect of the lymphocytes by Annexin V/PI

To investigate the cell viability effect of exosomes on lymphocytes, Annexin V-/PI-ratio was performed and analyzed (Fig. 4A). The combination of Annexin with PI allows discrimination between live cells (Annexin –, PI –), early apoptotic (Annexin –, PI +), late apoptotic (Annexin +, PI –) cells, and necrotic cell (Annexin +, PI+). Samples were acquired on a CytoFLEX, and data were analyzed with FlowJo. Cell survival ratio of peripheral blood mononuclear cells (PBMC) significantly decreased with IFN+Exo in patients with BU compared to both control BU and IFN-Exo groups (Fig. 4B).

**Figure 4.**
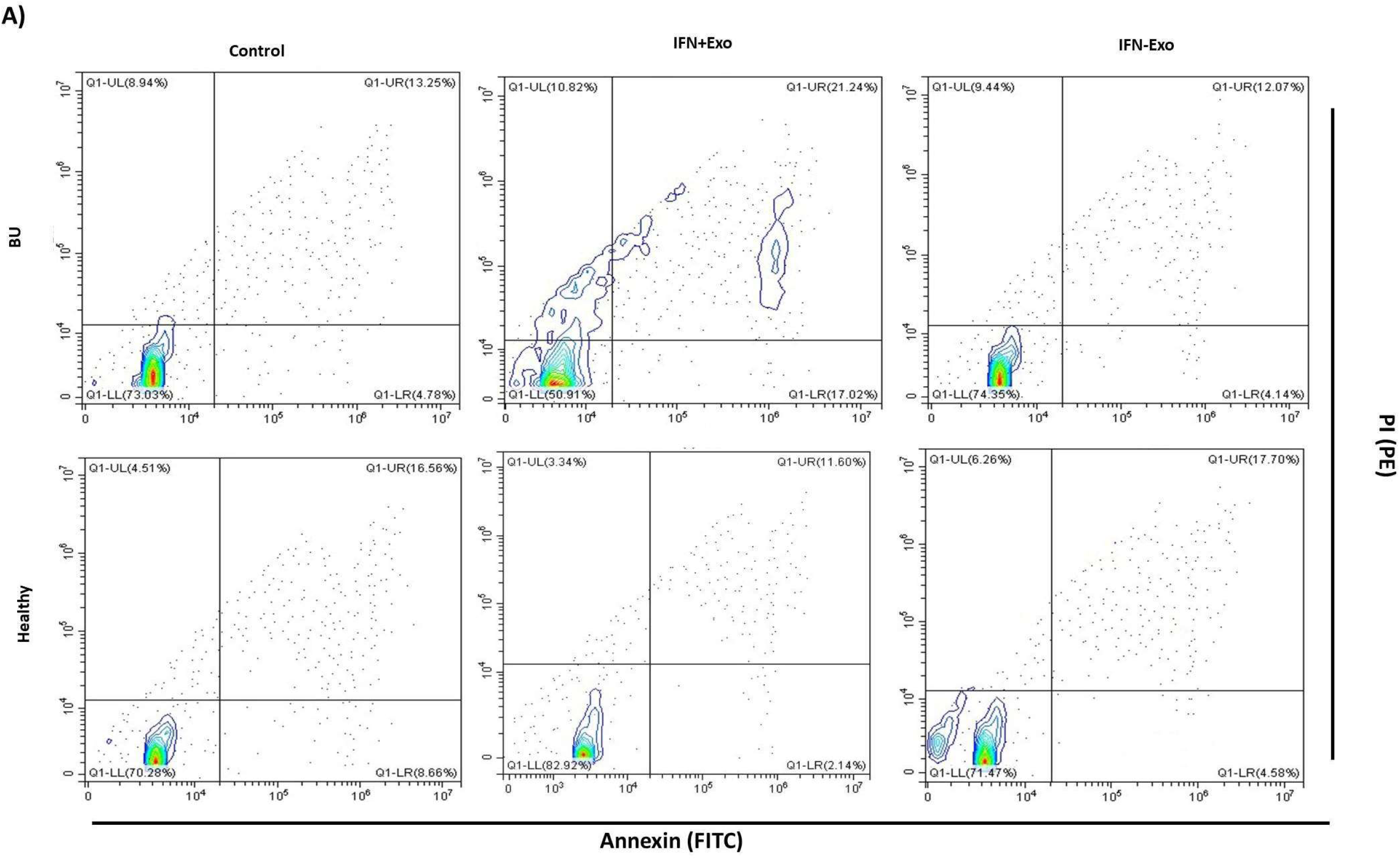
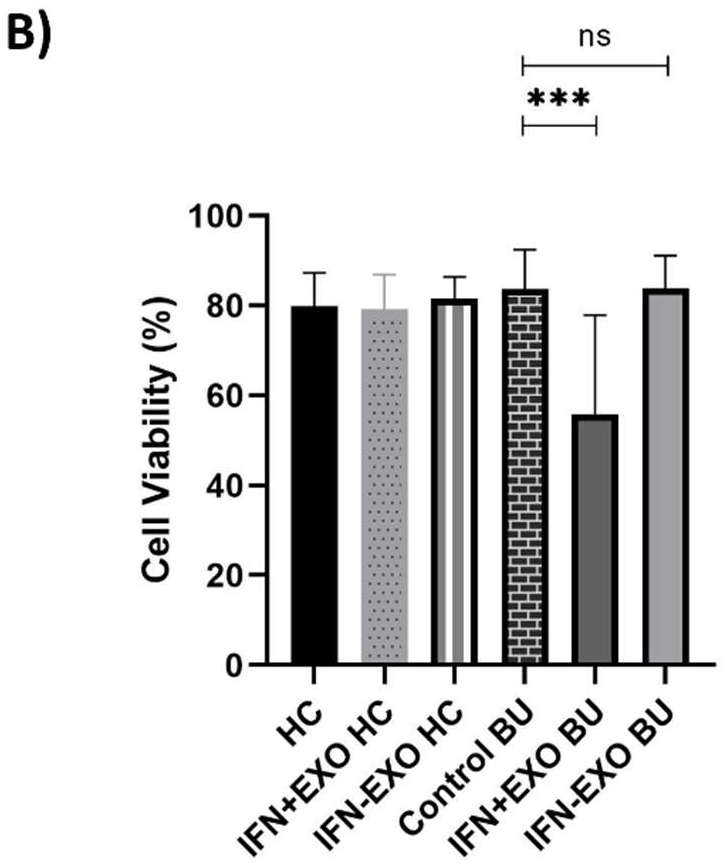
Cell viability analysis of lymphocytes. Lymphocytes were stained with Annexin V (FITC) and PI (PE) to analyze cell survival of lymphocytes and **(A)** displayed by flow cytometry. **(B)** In statistical analysis, cell survival ratio of PBMC decreased with IFN+ Exo in patients with BU compared to control BU, and it is statistically significant (n=15) (ns; P>0.05, ***P < 0.001). Results are shown as mean ± SD. *Patients with BD uveitis (BU), Exosomes of IFN-γ primed (IFN+Exo) and non-primed (IFN-Exo) AdMSCs, Healthy Controls (HC)*.

### 3.5 Determination of Fas/Fas Ligand pathway

Both lymphocytes were cultured in the presence and absence of IFN-γ exosomes. Fas/FasL ratio was determined for apoptosis after 3 days of culture, and changes in Fas/FasL expression on PBMC were examined between different groups (Fig. 5A). Fas/FasL apoptotic signaling of PBMC significantly increased with IFN+Exo stimulation in patients with BU compared to the control BU and IFN-Exo groups (Fig. 5B).

**Figure 5.**
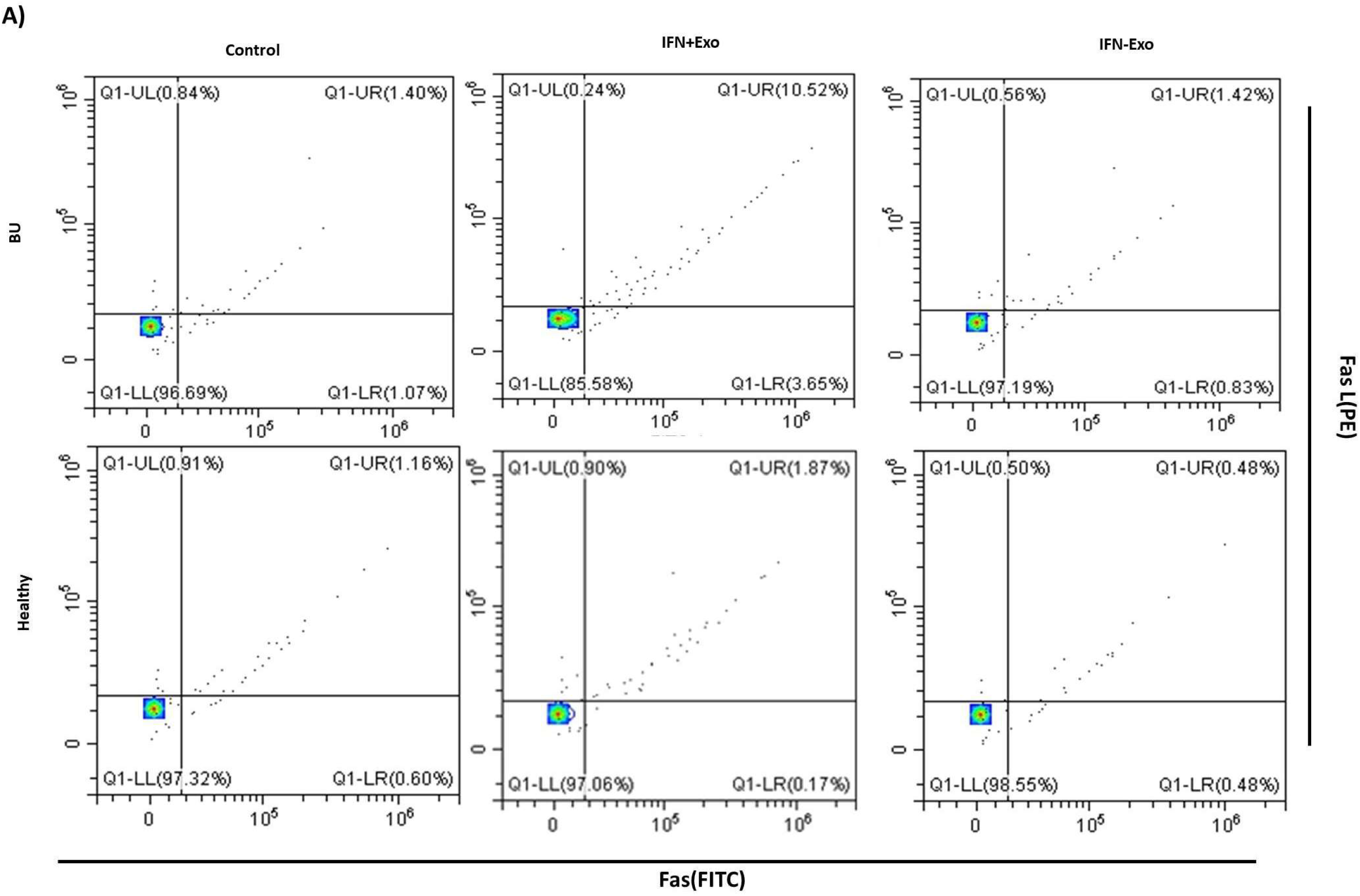
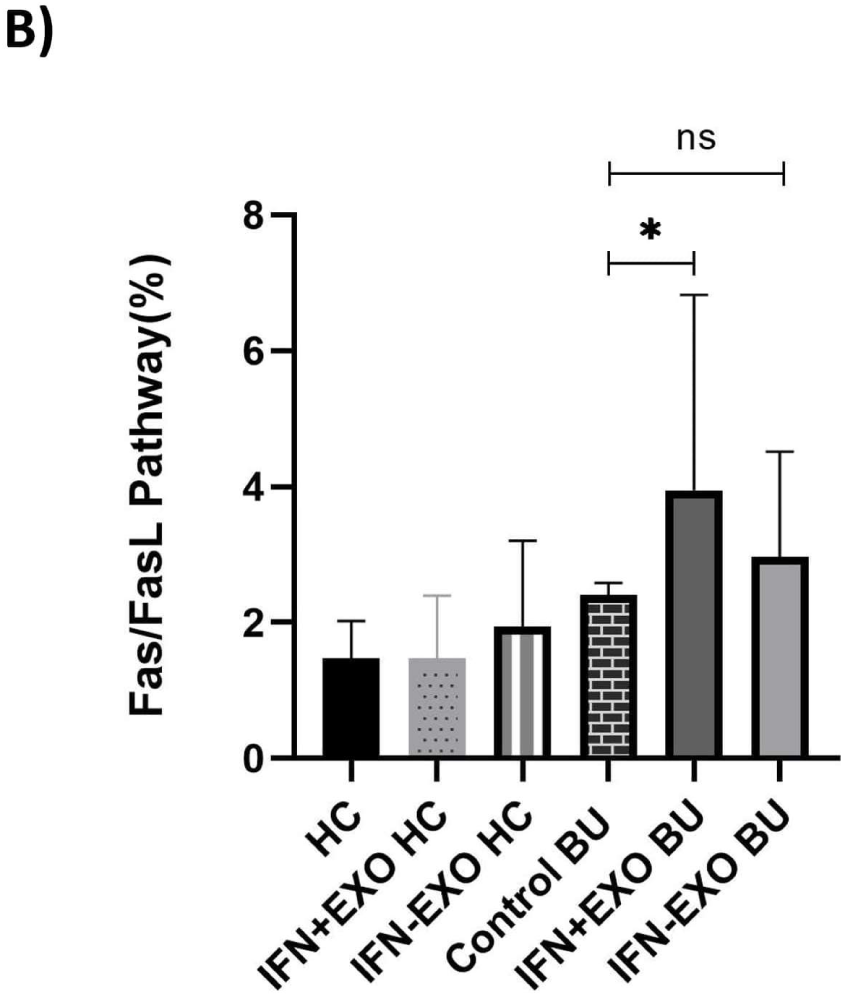
Evaluation of Fas and FasL Ratio. **(A)** After 72h of culture, the effect of IFN+ Exo on the proliferation of lymphocytes was demonstrated by flow cytometry. Lymphocytes were stained with anti-human CD178 (Fas-L) and anti-human CD95 (Fas) in order to analyze the ratio of Fas/FasL in PBMC of patients with BU (n=15). **(B)** The apoptotic pathway of PBMC increased with IFN+Exo in patients with BU compared to the control BU. (*P < 0.05). * denotes statistical significance compared to control BU. (#P<0.05). # denotes statistical significance compared between the IFN+/−Exo BU. Results are shown as mean ± SD. *Patients with BD uveitis (BU), Exosomes of IFN-γ primed (IFN+Exo) and non-primed (IFN-Exo) AdMSCs, Healthy Controls (HC)*.

### 3.6 Inflammatory Response

To assess the immunomodulatory effects of exosomes on lymphocytes, we evaluated pro-inflammatory and anti-inflammatory cytokine levels in PBMC of patients with BU and healthy controls. After 72h of culture, IFN-γ, TGF-β, and IL-17 were high in PBMC cultures of patients with BU compared to healthy controls. When PBMCs were treated with IFN+Exo, significantly decreased levels in IL-17 and TGF-β and significantly increased levels of IL-10 compared to patients with BU (Fig. 6). The results revealed that cytokines of interest did not alter after both exosome treatments in healthy controls (Figure S2.). In the current study, cytokines expressions of active and inactive patients with BU were evaluated by RT-PCR (Figure S1.). It was revealed that there was a significant difference in proinflammatory cytokine levels between active patients with BU when compared to both inactive BU and the healthy control groups. As no active inflammation was detected in inactive patients with BU, all experiments were performed with patients who had active BU.

**Figure 6.**
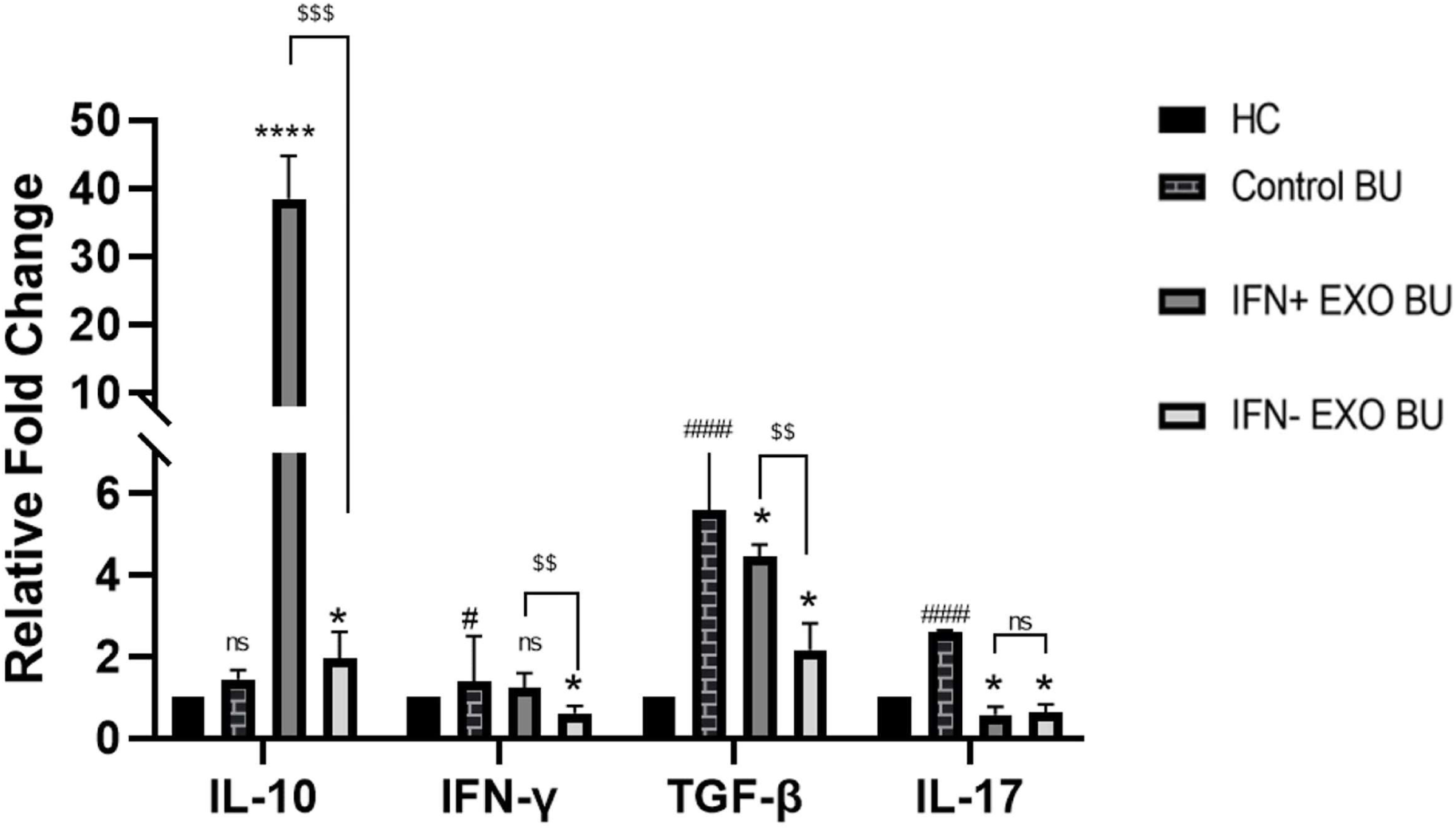
Cytokine levels in culture supernatants. After 72h of culture, levels of IFN-γ, TGF-β, and IL-17 were significantly higher in PBMC cultures of patients with BU compared to healthy controls (#: P<0.05, ####: P<0.0001; denotes statistical significance between control BU and control HC groups). When PBMCs were treated with IFN+Exo or IFN-Exo, levels of all pro-inflammatory cytokines decreased significantly except IFN-γ levels in IFN+Exo group (*: P<0.05, ****P<0.0001; denotes statistical significance between treatment groups and control BU).;Increase in IL-10 levels were significantly higher with IFN+Exo treatment whereas IFN-Exo treatment decreased levels of proinflammatory cytokine levels significantly when compared to IFN+Exo group ($$:P<0.01, $$$:P<0.001 denotes statistical significance between treatment groups). Results are shown as mean ± SD. *Patients with BD uveitis (BU), Exosomes of IFN-γ primed (IFN+Exo) and non-primed (IFN-Exo) AdMSCs, Healthy Controls (HC)*.

## 4 Discussion

An in-vitro cell culture model was employed to assess the impact of exosomes secreted from IFN-γ-primed and non-primed AdMSCs on the PBMC of patients with BU. This is the first study to investigate the effects of AdMSC-derived exosomes on PBMC-related immune response in patients with BU. MSCs activated with IFN-γ have a robust immunomodulatory effect due to the immunoactive factors upregulation [23]. The current study also demonstrated the differences between primed and non-primed MSC-derived exosomes on the BU lymphocytes. The study’s results revealed that IFN+Exo decreased the viability and increased apoptosis pathways of T lymphocytes in BU. Exosomes raised IL-10 levels as an anti-inflammatory cytokine and decreased IL-17, TGF-β and IFN-γ levels as a pro-inflammatory cytokine in patients with BU’s lymphocytes. Nevertheless, AdMSC-derived exosomes did not affect on T lymphocyte proliferation.

In the present study, we employed exosomes of AdMSCs as the primary treatment options rather than using AdMSCs themselves. The injection of MSCs have significant limitations, such as the limited maximum quantity of MSCs injected and the suspension volume. Exosomes of MSCs hold a particular potential to be used as therapeutic tools for resolution of inflammatory conditions including BD as they are not prone to disadvantages of cellular therapies. For instance, exosomes can pass through biological barriers like the blood-brain-barrier [24]. They may modulate the immune system through multiple mechanisms without having the possible concerns that can be encountered with MSCs due to undesired cell differentiation [25]. The possible reinfusion of unpurged autoreactive lymphocytes during stem cell rescue procedures with possible acute graft-versus-host disease, and the need for continuous immunosuppression following the treatment are two other significant limitations of stem cell rescue techniques [26]. From that aspect, one of the major advantages of exosomes over MSCs, is their hypo-immunogenicity. The absence of the major histocompatibility complex (MHC)-II reduces the possibility that exosomes will be identified by immune system which, prevents an increase immune response. Due to the low-level expression of MHC-I on exosomes, they are less likely to elicit cytotoxic T-cell responses. They reduce the risk of triggering immune reactions, which can be advantageous in therapeutic applications where immune modulation is preferred, such as treating autoimmune diseases and other inflammation diseases. Immune response dysregulation is a common feature of autoimmune diseases. [19]. By delivering therapeutic reagents to immune cells, such as antigens, RNA, or regulatory proteins, exosomes may be able to target MHC molecules, receptors involved in immune cell activation, cytokines, and signaling molecules, challenging autoimmune reactions.

In the current study, we did not see any difference in cell proliferation in the treatment groups (Fig. 3). Despite the culture duration is limited for demonstrating any effect on the proliferation of PBMC, our study’s results are consistent with previous reports which no difference in cell proliferation between patients with BU and healthy controls [27]. Moreover, exosomes derived from late passages of MSCs do not effectively suppress PBMC proliferation. It suggests that the anti-inflammatory phenotype of MSCs was attenuated during extensive subculturing [28]. Similarly, exosomes from late-passage stem cells in our study may not affect on proliferation. The exosome treatments proposed do not imply any immunosuppression targeting mononuclear cell proliferation, which could potentially increase the safety profile of the treatment strategy.

The effects of exosomes on the cell survival of T lymphocytes of patients with BU were investigated (Fig. 4). It is remarkable that the cell viability reduction effect of IFN+Exo treatment was limited to BU group and no significant impact over the healthy control group was observed. This limited activity could be related to Fas-FasL-mediated apoptosis, which is a mechanism for preserving immunological homeostasis by eliminating autoreactive cells during the maturation of the immune system [29, 30]. Exosomes induce apoptosis of T lymphocytes through the interaction of FasL and Fas [31–33]. For example, exosome injection treated established collagen-induced arthritis and decreased the inflammatory response by modulating T cell activity in vivo through an MHC class-II and partially Fas ligand/Fas-dependent pathway [34]. Another study showed that myeloid-derived suppressor cells (MDSCs)-derived exosomes precipitate T-cell apoptosis through hyper-activation or repeated stimulation resulting in increased levels of ROS production and activation of the Fas/FasL pathway [35]. A significant increase in the Fas-FasL ratio was observed in our study (Fig.5). It supports the role of Fas-FasL-mediated apoptosis in maintaining immunological homeostasis and eliminating autoreactive cells during immune system maturation.

Studies suggest that the pathogenesis of BD is primarily driven by an immune response of Th1 and Th17 cells [7, 36, 37]. Notably, elevated levels of cytokines associated with Th1/Th17 cells, such as IL-2, IFN-γ, IL-6, IL-12, IL-17, and IL-23, have been documented in individuals with BD [8]. Thereby, the present study focused mainly on the modulation of Th1, Th17, and Th2 cells as well as related cytokines after 72 hours of treatment with IFN-γ-stimulated AdMSC-derived exosomes. Nojehdehi et al. [38] showed AdMSC-derived exosomes decreased IL-17 levels in autoimmune diabetes. The discovery of the role of TGF-β in Th17 cell activation once again revealed the pro-inflammatory effect of TGF-β [39]. Studies suggest TGF-β is required for cell differentiation of Th17 plays crucial roles in the pathogenesis of several autoimmune diseases such as rheumatoid arthritis and inflammatory bowel disease [39, 40]. In our study, the levels of TGF-β and IL-17 were significantly reduced in both the IFN+Exo and IFN-Exo treated BU groups when compared to the control BU (Fig.6). It is noted that the decrease of TGF-β in the IFN-Exo group was even significantly higher than the IFN+Exo group, while no significant difference was found between the groups on the reduction of IL-17. However, the IFN+Exo group had no effect in reduction of IFN-γ whereas the IFN-Exo group resulted in a significant decrease. These data indicated that the decrease in pro-inflammatory cytokines was not associated with exosomes of primed-AdMSCs. The pro-inflammatory cytokines are inhibited by the anti-inflammatory cytokine IL-10, which plays a major role in the pathogenesis of BD uveitis [41]. Li et al.[42], found that human umbilical cord mesenchymal stem cell (hUMSC)-exosomes suppressed TNF-α and IL-1β secretion and increased IL-10 secretion. Similarly, IL-10 expression increased in the MSC-Exo-treated mice with myocardial I/R injury [43]. In this study, we demonstrated that IL-10 gene expression was significantly elevated in both IFN+Exo and IFN-Exo treated BU groups compared to the control BU (Fig. 6). Remarkably, the nearly 40-fold increase of IL-10 in the IFN+Exo group could be considered as the main anti-inflammatory pathway triggered by the AdMSCs primed with IFN-γ-derived exosomes. They can potential to increase the effect of anti-inflammatory macrophages that are responsible for tissue repair, including the reduction of fibrosis and stimulation of tissue regeneration [44]. Given that the expansion of IL-10 downregulates Th17 function [45], the resulting immunomodulation would be able to resolve inflammatory response related to BU. Hence, exosomes may be an effective therapy option for uveitis with Behçet’s disease, particularly for patients who do not respond to drugs. However, further research and comparisons between the exosomes and drugs are required.Their potential clinical benefits could include better therapeutic outcomes and personalized treatment alternatives, opening the path for more effective and less invasive approaches.

The current study has some limitations. We were able to evaluate limited number of immune response markers. While the cytokines examined in this study have a significant impact on the BU, for a clearer understanding of the treatment of BU, further studies of other major cytokines that are found to be responsible for BD pathogenesis should be investigated in larger cohorts. In particular, IL-1β and TNF-α as pro-inflammatory agents are important in the development of BD [46, 47]. Addressing these limitations in further investigations will be essential for advancing our understanding of exosome treatment in BU. The effect of intersex immune response variations and the influence of genetic factors over cytokine production should also be considered to clarify further the mechanisms [48, 49]. Alternatively, the enhanced anti-inflammatory effects of IFN+Exo and IFN-Exo should be supported by enzyme-linked immunosorbent assay (ELISA) analysis at the protein level. It provides more information on immune response after exosomes, supporting the results of RT-PCR and expanding our knowledge of the exosome treatments’ overall therapeutic impact.

## Conclusion

Our study demonstrated the immunomodulatory effects of adipose-derived mesenchymal stem cell exosomes over inflammatory cells isolated from subjects with BU. Once primed with IFN-γ, these exosomes could repress excessive T-cell activation by modulating cytokine secretion in favor of anti-inflammatory pathways. Exosomes of AdMSCs are distinguished as a promising and innovative treatment approach in BU.

## Supporting information

Supplemental Datas

## Acknowledgments

The authors would like to thank TUBITAK for the financial support. The authors also gratefully acknowledge the use of the services and facilities of the Koç University Research Center for Translational Medicine (KUTTAM) and Koç University Surface Science and Technology Center (KUYTAM), funded by the Presidency of Turkey, Head of Strategy and Budget. We thank the participants of the study.

