## Supplemental Datas for "Effects of Primed Adipose Mesenchymal Stem Cell-Derived Exosomes on Immunomodulation in Behcet Uveitis"

**Supplementary materials**

| Genes | Primer Sequences (5’ to 3’) | |
| --- | --- | --- |
| TGF-β1 | F | GGC CAG ATC CTG TCC AAG C |
|  | R | GTG GGT TTC CAC CAT TAG CAC |
| IFN-Ɣ | F | TCG GTA ACT GAC TTG AAT GTC CA |
|  | R | TCG CTT CCC TGT TTT AGC TGC |
| IL-17 | F | CCTA CAA TGC TGA CAA CCGC |
|  | R | GGG CTG TGT AGA GGG CAA TC |
| IL-10 | F | GAC TTT AAG GGT TAC CTG GGT TGT |
|  | R | TCA CAT GCG CCT TGA TGT CTG |
| GAPDH | F | ATG GAA ATC CCA TCA CCA TCT T |
|  | R | CGC CCC ACT TGA TTT TGG |

**Table S1:** List of primer sequences for RT-PCR

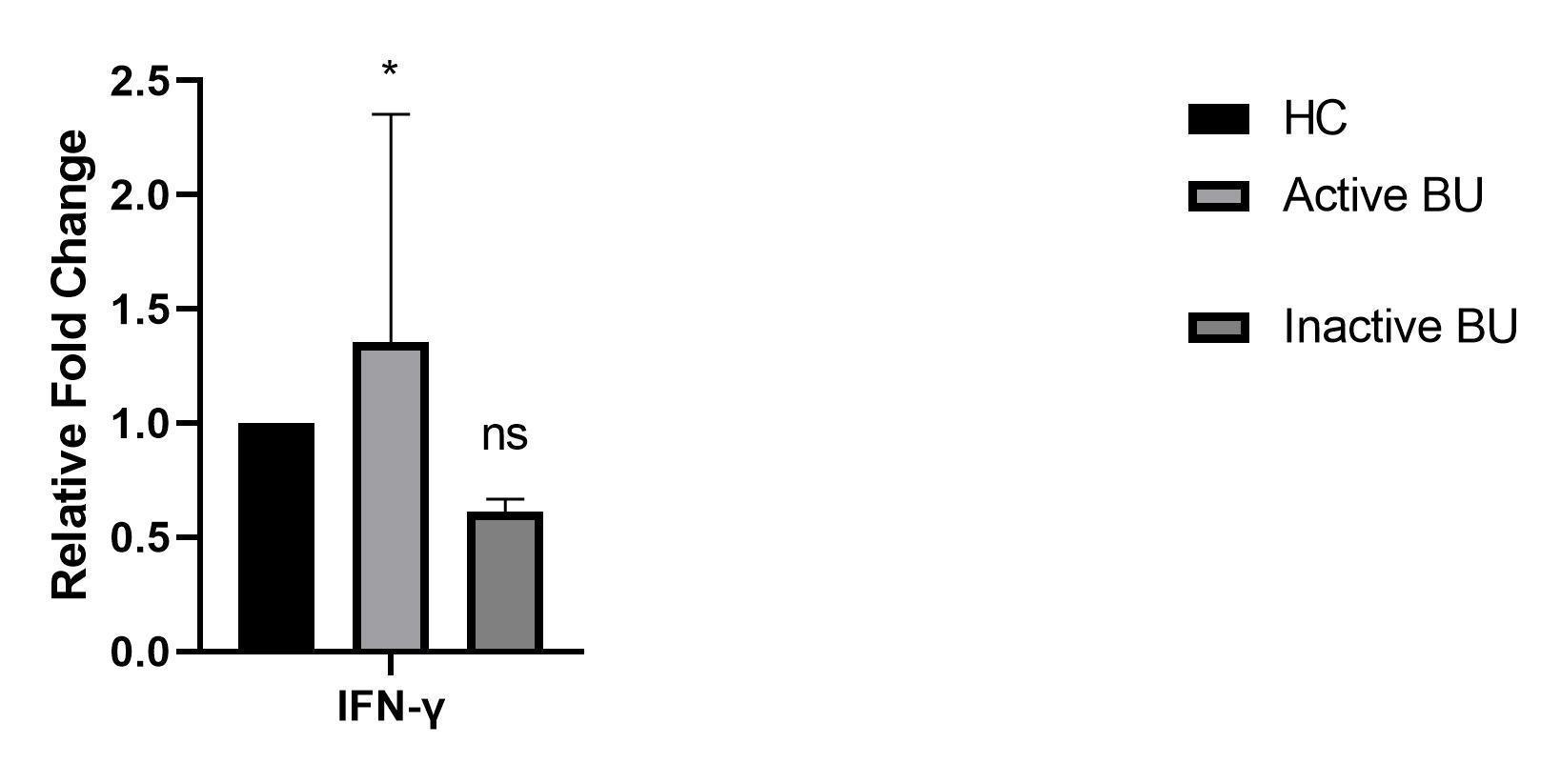

**Figure S1. Comparison of Proinflammatory Levels in Patients of Active and Inactive Patients with BU.** IFN-γ level was high in PBMC cultures of Active patients with BU compared to healthy individuals. *P <0.05. IFN-γ level did not change in PBMC cultures of Inactive patients with BU compared to healthy control. P>0.05. Results are shown as mean ± SD. *Patients with BD uveitis (BU), Exosomes of IFN-γ primed (IFN+Exo) and non-primed (IFN-Exo) AdMSCs, Healthy Controls (HC).*

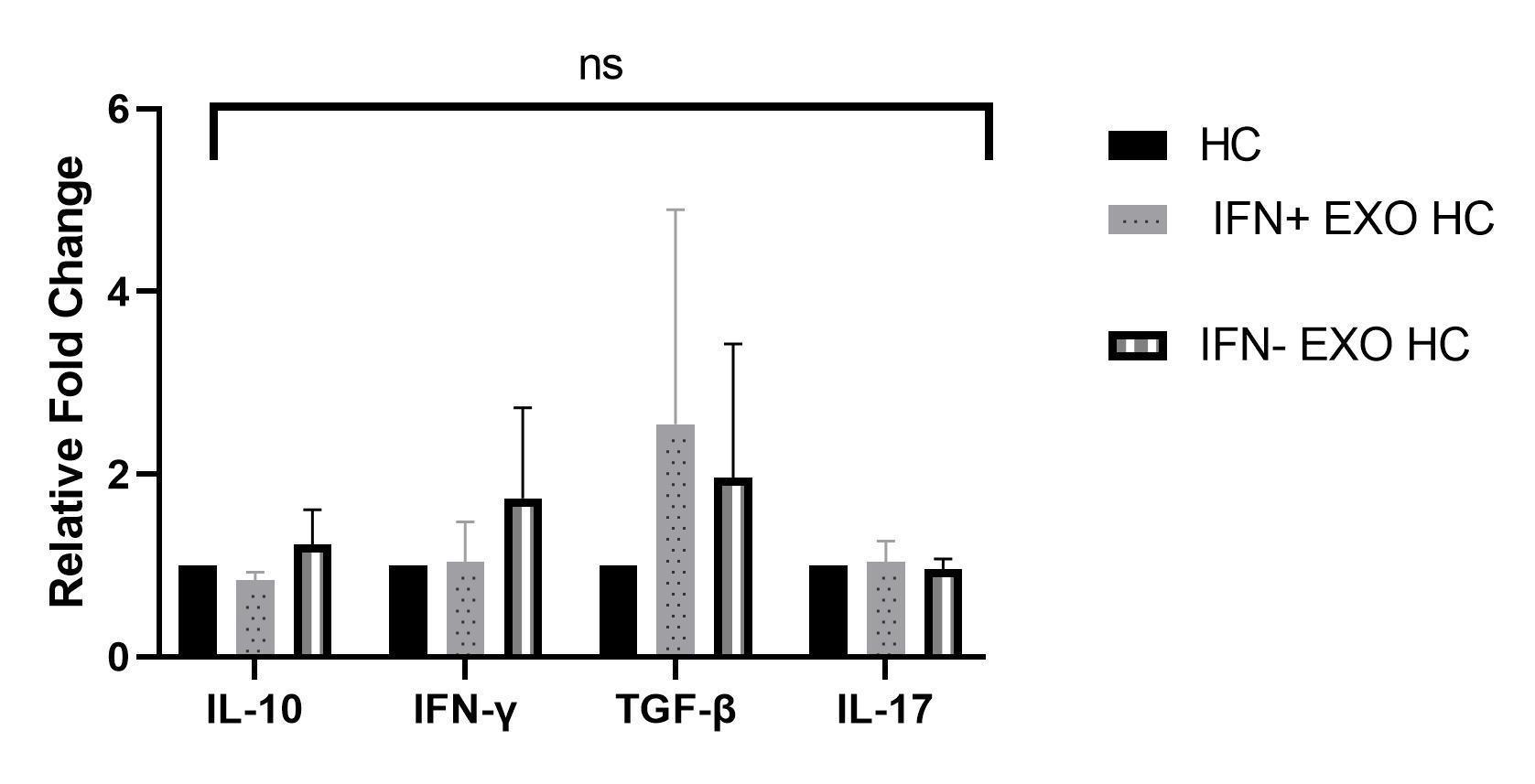

**Figure S2. Cytokine levels in culture supernatants of healthy individuals**. After 72 h of culture, exosome treatments do not affect healthy controls. P>0.05. Results are shown as mean ± SD. *Patients with BD uveitis (BU), Exosomes of IFN-γ primed (IFN+Exo) and non-primed (IFN-Exo) AdMSCs, Healthy Controls (HC).*
